# Maintaining robust size across environmental conditions through plastic brain growth dynamics

**DOI:** 10.1101/2020.09.01.277046

**Authors:** Ansa E. Cobham, Brent Neumann, Christen K. Mirth

## Abstract

Organ growth is tightly regulated across environmental conditions to generate appropriate final size. While the size of some organs is free to vary, others need to maintain constant size to function properly. This poses a unique problem: how is robust final size achieved when environmental conditions can alter some major growth processes? While we know that brain growth is “spared” from the effects of the environment from humans to fruit flies, we do not understand how this process alters growth dynamics across brain compartments. Here, we explore how this robustness in brain size is achieved by examining differences in growth patterns between the larval body, the brain, and a brain compartment – the mushroom bodies – in *Drosophila melanogaster* across both thermal and nutritional conditions. We identify key differences in patterns of growth between the whole brain and mushroom bodies that are likely to underlie robustness of final organ shape. Further, we show that these differences produce distinct brain shapes across environments.

**Significance of Study:** A long-standing question in Biology has been how fully functional multicellular organisms with highly specialized organs are generated, given that organs initiate growth at different times across development. Although the genetic mechanisms that underlie growth has been studied extensively, we are yet to understand how growth pattern of organs produces distinct final shapes across changing environmental conditions. We use the Drosophila brain, to reveal that key differences in growth dynamics are likely to underlie robustness of final organ shape and are tuned by nutrition and temperature. Further deepening our knowledge of how final organ shape is maintained across environmental conditions.

## Introduction

How are the shapes and sizes of growing organs regulated throughout development to generate a fully functional multicellular animal with highly specialized parts? This seems particularly difficult to understand given that body parts initiate growth at different times, and further grow at different rates and with differing dynamics (Andersen et al., 2013; Eder et al., 2017; Huxley, 1932). While some organs show exquisite sensitivity to environmental conditions, known as plasticity, changing their shape and size with changes in nutrition, temperature, and other conditions (Bateson, 2017); other organs maintain relatively constant final sizes across conditions (Bateson, 2017; Nijhout, 2002). The properties that allow growth to resist perturbations in environmental conditions contribute to robustness in development (Bateson, 2017; Mirth & Shingleton, 2019; Nijhout, 2002). As organs vary in sensitivity to environmental perturbations, animals that develop in different environments will differ in their body size and shape (Mirth & Shingleton, 2012). Understanding the properties of organ growth that allow them to be either plastic or robust to environmental conditions is key to uncovering how correct, functional body form is achieved

Extensive studies in insects have described how the patterns of growth across organs generate variation in size and shape of the adult body (Andersen et al., 2013; Mirth & Shingleton, 2012, 2019; Nijhout et al., 2014). Varying growth dynamics can occur either at the level of an individual organ or through coordinating growth processes among organs relative to the growing body (Huxley, 1932; Shingleton & Frankino, 2018). Also, environmental conditions can act to alter each of these growth properties (Miner et al., 2000; Nijhout & Grunert, 2010; Shingleton et al., 2009; Shingleton et al., 2008).

Across a wide variety of animals, including mammals and insects, the brain is generally less sensitive to changes in environmental conditions than other organs of the body (Cusick & Georgieff, 2016). This is commonly referred to as brain sparing (Cohen et al., 2015). In humans, newborns raised under reduced nutrient availability or oxygen supply have reduced weight and body sizes, and disproportionately large heads (Cohen et al., 2015; Cox & Marton, 2009). Illustrating that the brain has built-in mechanisms to ensure its size is not compromised.

Brain differentiation in *Drosophila*, occurs in the embryo, a stage that is protected from nutrient restriction as the embryos are not fed. However, most brain growth occurs during the larval stages, and nutrition plays an important role (Yuan et al., 2020). Poor nutrition, especially in the later stages of larval development, produces small sized adults (Mirth & Shingleton, 2012), but with proportionally larger brains than those reared under nutrient rich conditions (Cheng et al., 2011). Brain growth is spared against poor nutrition via the action of the glial secreted tyrosine kinase-like insulin receptor called Alk and its ligand Jelly Belly (Jeb) (Cheng et al., 2011). Alk activates downstream effectors of the insulin signalling pathway and downstream targets of the Target of Rapamycin (TOR) kinase bypassing amino-acid sensing in the absence of nutrient cues, to ensure that the size and composition of cells in the brain is maintained even when larvae are starved (Cheng et al., 2011; Lanet & Maurange, 2014).

While these findings highlight a genetic mechanism through which brain sparing occurs, they do not explain how brain growth adjusts with extended larval growth periods caused by poor nutrition. If Alk signalling maintains high growth rates in starved larvae as it does in fed, the extension of developmental time caused by starvation would cause brains to overgrow. But since this does not happen, it suggests that the growth dynamics in brains of starved larvae adjust to avoid overshooting their size with longer growth periods.

This could happen in several ways (Fig 1). Firstly, when larvae are starved, they could maintain constant growth rates within the brain, but then stop growing once a target size is reached (Figure 1A). This would result in a growth trajectory that reaches an asymptote. Alternatively, starved larvae could delay the time at which they initiate brain growth, but once initiated, they maintain constant growth rates (Figure 1A). Larvae raised under different nutritional conditions would show exponential growth trajectories, where exponential growth would be initiated at different times depending on the rearing conditions of the larvae (lagged exponential model). Finally, Jeb and Alk might not act to ensure insulin and TOR signalling are maintained at constant levels. Instead, these pathways might tune both growth rates and the timing at which growth is initiated, to adjust for the extended growth period (Figure 1A). This would result in changes in both the time at with growth was initiated as well as the growth rate. These differences in growth dynamics are important, as each implies a different mechanism for adjusting brain size with environmental conditions.

**Figure 1:**
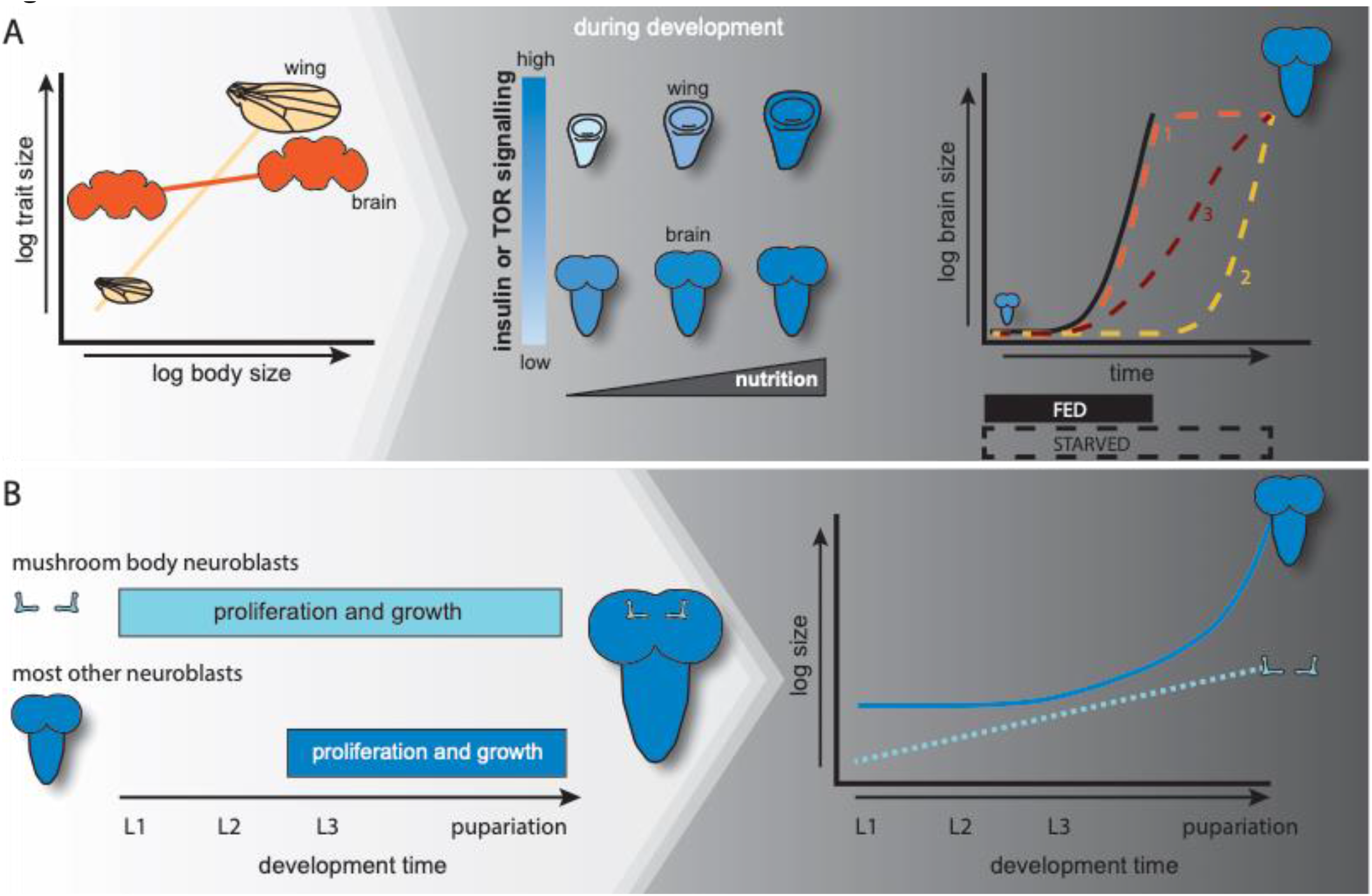
How do the growth dynamics of the whole brain and the mushroom bodies vary. (A) Hypothesis 1: Mushroom bodies proliferate throughout larval development, while most of other neuroblasts in the brain remain quiescent and reinitiate proliferation in the late second instar (L2). These differences in proliferation could results in differences in the dynamics of mushroom body growth when compared to the whole brain. While we expect that the whole brain would show a lag period where it does not growth, followed by a period of exponential growth, the mushroom body might show constant (linear) increases in size across the larval stages of development (dashed line 1). Alternatively, the mushroom body might show similar growth dynamics, with shallower increase in growth rate in later development (dashed line 2). Differences in growth dynamics between the mushroom bodies and the whole brain would suggest that they are regulated in distinct manners under changing environmental conditions. (B) In comparison to other organs like the wing, adult brain size changes little with changes in body size. The reason that this is thought to occur is that insulin and target of rapamycin (TOR) signalling is kept high in the brain even under poor nutrient conditions via the action of Jeb/Alk. High levels of insulin or TOR signalling would suggest that brains would maintain constant growth rates even across environmental conditions – like starvation – that induce prolonged larval development. To maintain constant size, this would mean that the brain would either need to grow at constant rates until it reached its target size and then stop (orange dashed line 1), or else delay the onset of growth until later (yellow dashed line 2). Alternatively, Jeb/Alk could tune insulin or TOR signalling levels such that the rate of growth was reduced to compensate for the extended development time (red dashed line 3).

Compared to the overall, different neuronal subclasses vary their rates of cell division in response to nutrition and other environmental conditions like temperature, light, and population densities during larval stages of growth (Heisenberg et al., 1995; Lin et al., 2013; Prokop & Technau, 1994; Wang et al., 2018). As most neuroblast populations enter quiescence in the early larval stages, the neuroblasts that give rise to the mushroom bodies – the paired neuronal structures important for olfactory processing and learning – continue to divide and differentiate from the first instar (L1) stage onwards. (Kunz et al., 2012). When faced with extremely poor nutritional conditions that reduce larval growth, the mushroom body neurons maintain division of the same neuronal cell types without differentiating (Lin et al., 2013; Rossi et al., 2017). In contrast to the mushroom bodies, the optic lobe neurons, which receive sensory input from the visual system, are only activated late in larval development and are highly sensitive to changes in nutritional environment, with the size of the neuron pool involved in initial proliferation highly dependent on nutrient availability (Lanet & Maurange, 2014). This suggests that specific brain regions differ in how they protect the whole brain from environmental perturbations.

These findings allow us to further propose a model of how the mushroom body compartments of the brain might maintain constant size in the face of changing environmental conditions. Firstly, because neuroblasts such as those that give rise to the mushroom body neurons begin proliferating much earlier than the majority of the brain neuroblasts, and proliferate throughout the larval instars, we might expect the size of these structures to increase constantly, or linearly, throughout larval development (Figure 1B). Most of the remaining neuroblasts of the brain initiate proliferation late in the second instar. Thus, we would expect a period of little or no discernible growth across the whole brain in the first two instars, punctuated by a rapid onset of growth in later development that would be best characterized by exponential growth with a time lag to its onset (Figure 1B). This would mean that growth dynamics in the mushroom body could differ significantly from that of the whole brain. Differences in their dynamics could indicate that growth is mediated by differing mechanisms between compartments, potentially dictating their response to environmental cues.

In the current study, we aim to determine how brain growth dynamics are regulated to ensure robust size across different environmental conditions, and whether all compartments of the brain regulate these dynamics in the same manner. To address this, we compared the growth patterns of whole brains and mushroom bodies, relative to the larval body, under standard rearing conditions. We then used altered nutritional and thermal conditions to explore how the dynamics of brain growth respond to environmental change. These studies reveal differences in the way the mushroom body compartment regulates its growth when compared to the whole brain, and highlight how growth dynamics are tuned by nutrition and temperature. With these studies, we deepen our understanding of how different brain regions maintain robustness across environmental conditions.

## RESULTS

### Comparing the growth dynamics of the larval body, whole brain, and mushroom bodies across larval development

Given that the mushroom body neuroblasts show different patterns of growth to the majority of other neuroblasts in the brain, our first goal was to devise methods to compare mushroom body growth to whole brain and larval body growth across all three larval instars. To ensure that we compared the growth of the same structures across developmental time, we required a marker that would be expressed throughout all three instars. Using the expression data available from the Janelia FlyLight project (http://flweb.janelia.org/cgi-bin/flew.cgi), we found that the GMR38E10 GAL4 line drove GFP expression in the vertical and medial lobes of the mushroom body neurons from hatch through to pupariation (Supplementary Figure 1). In the late L3 stage, GFP expression was not apparent in the mushroom body calyx (Supplementary Figure 1), which is the dendritic projections of Kenyon cell bodies (Supplementary Figure 2a and 2b). Thus, to be able to compare measurements across all stages of development we excluded the calyx and peduncles from our analyses and measured only the ventral and medial lobes for mushroom body volume (Supplementary Figure 2a and 2b).

We next sought to compare the dynamics of larval, whole brain, and mushroom body growth. Log-transformed larval growth increased steadily throughout the first, second, and third instar stages (Supplementary Figure 3A-C, Supplementary Table 1). Linear models explain 68%, 55%, and 78% of the variation in larval volume over time for L1, L2, and L3 respectively (Supplementary Table 1, adjusted R^2^ values). Similarly, the mushroom body displayed steady linear growth throughout all three instars (Supplementary Figure 3G-I, Supplementary Table 1), with linear models explaining 43%, 55%, and 77% of the variance in mushroom body volume over time for the L1, L2, and L3 respectively (Supplementary Table 1, adjusted R^2^ values). In contrast, for whole brain volume we observed a slight, but significant, decrease in whole brain volume with time in the L1 (Supplementary Figure 3D, Supplementary Table 1). In this case, the linear model explained only 4% of the variance in whole brain volume in the L1 (Supplementary Table 1, adjusted R^2^ values). There was no significant change in brain volume with time across the L2 stage (Supplementary Figure 3E, Supplementary Table 1). In the L3, whole brain volume shows a non-linear relationship with time, curving upwards. This suggests that whole brain growth speeds up as the third instar progresses (Supplementary Figure 3F). Curiously, at 0 hours after the moult to both L2 and L3, brain volume appears to increase despite no evidence of positive growth during the L1 or L2 instars. We cannot tell whether this is a random sampling effect or if this results from a burst of growth during the moult cycle itself, which we could not accurately sample.

Our results thus far suggest that whole brain growth is regulated differently to that of the larval body and mushroom bodies. To formally test this, we fit our growth data with both linear models and a range of non-linear models commonly used to describe growth dynamics, including second order polynomial, exponential, lagged exponential, and power models (Karkach, 2006). Each of these models infers something different about growth. The second order polynomial model assumes that the organ will have periods where its growth increases steadily with time, as well as periods during which growth rates slow down; exponential models describe growth that speeds up exponentially over time; lagged exponential models are similar to exponential models, but infer a period of slow or no growth followed by a switch to exponential growth; and the power model implies that growth increases according to a power function. We assessed which model best fit our growth data for each trait using two different model selection methods: Akaike’s Information Criteria (AIC) and Bayesian Information Criteria (BIC), both of which estimate the quality of each model relative to the others, penalizing models with a higher number of parameters to avoid overfitting the data. The model with the lowest AIC and BIC values provides the best fit for the data. Where these values were close between models, we selected the simplest model (i.e. the model with the fewest parameters). We restricted these comparisons to L3 growth, since the whole brains did not show significant positive growth in the L1 and L2 stages.

For growth in the larval body and mushroom body, we found that linear models provided the best fit to our data (Supplementary Table 2). This means that the growth rates in the larval body and mushroom body do not change over time in the third instar. Whole brain growth, on the other hand, was best fit with a lagged exponential model. This indicates that in the early stages of the third instar the whole brain grew very slowly. After this initial lag phase, the rate of whole brain growth increased exponentially. Taken together, these data suggest that while the larval body and mushroom body growth rates do not change with time over the third instar, the whole brain undergoes a period of little growth, followed by a second phase of rapidly increased growth in the L3.

### Developmental time and growth dynamics are modulated by changes in nutrition and temperature

We next sought to determine how brain size remains robust when developmental time becomes extended as a result of altered environmental conditions. To do so, we first determined the diet and temperature conditions that produced the most differences in brain growth. We reared larvae on 5 different diets of 10%, 12.5%, 25%, 50%, and 100% and three temperatures 18ºC, 25ºC, or 29ºC. Our preliminary data showed that we could achieve the greatest range of effects by comparing the 10%, 25% and 100% diets and 25ºC and 29ºC rearing temperatures (Supplementary Figure 4). We compared growth rates in the L3 across these six environmental conditions. Changing the diet and/or rearing temperature altered the time it took for animals to initiate metamorphosis at pupariation (white pre-pupae). Compared to animals grown under standard conditions (25ºC and 100% food), animals reared on food with only 10% of the normal caloric content took the longest to pupariate (90 and 80 hours after the moult at 25ºC and 29ºC respectively, compared to 42 hours at 25ºC on 100% food). At 25ºC, pupariation was delayed to 50 hours after the moult when larvae were reared on 25% food. Development time was similar between the 25% and 100% food conditions at 29ºC (42 hours from moult to white pre-pupae).

Given these differences in development time across nutritional and thermal conditions, we next defined how this changed growth dynamics of the mushroom body, whole brain, and larval body. For each condition, we sampled 5-7 time points across the L3 stage, with the last 2 time points corresponding to the wandering and white prepupal stages, respectively. Diluting the food reduced growth rates of the larval body at both temperatures (Figure 2A, B, Supplementary Table 3). Overall, the larval body grew more slowly when larvae were reared at 29ºC compared to 25ºC (Figure 2A, B, Supplementary Table 3). Larvae grew slowest on 10% food at 29ºC and fastest on 100% food at 25ºC (Figure 2A, B, Supplementary Table 3), resulting in a significant interaction between time, food, and temperature. These data provide a convenient proof-of-principle that we can alter growth dynamics by manipulating food and temperature.

**Figure 2:**
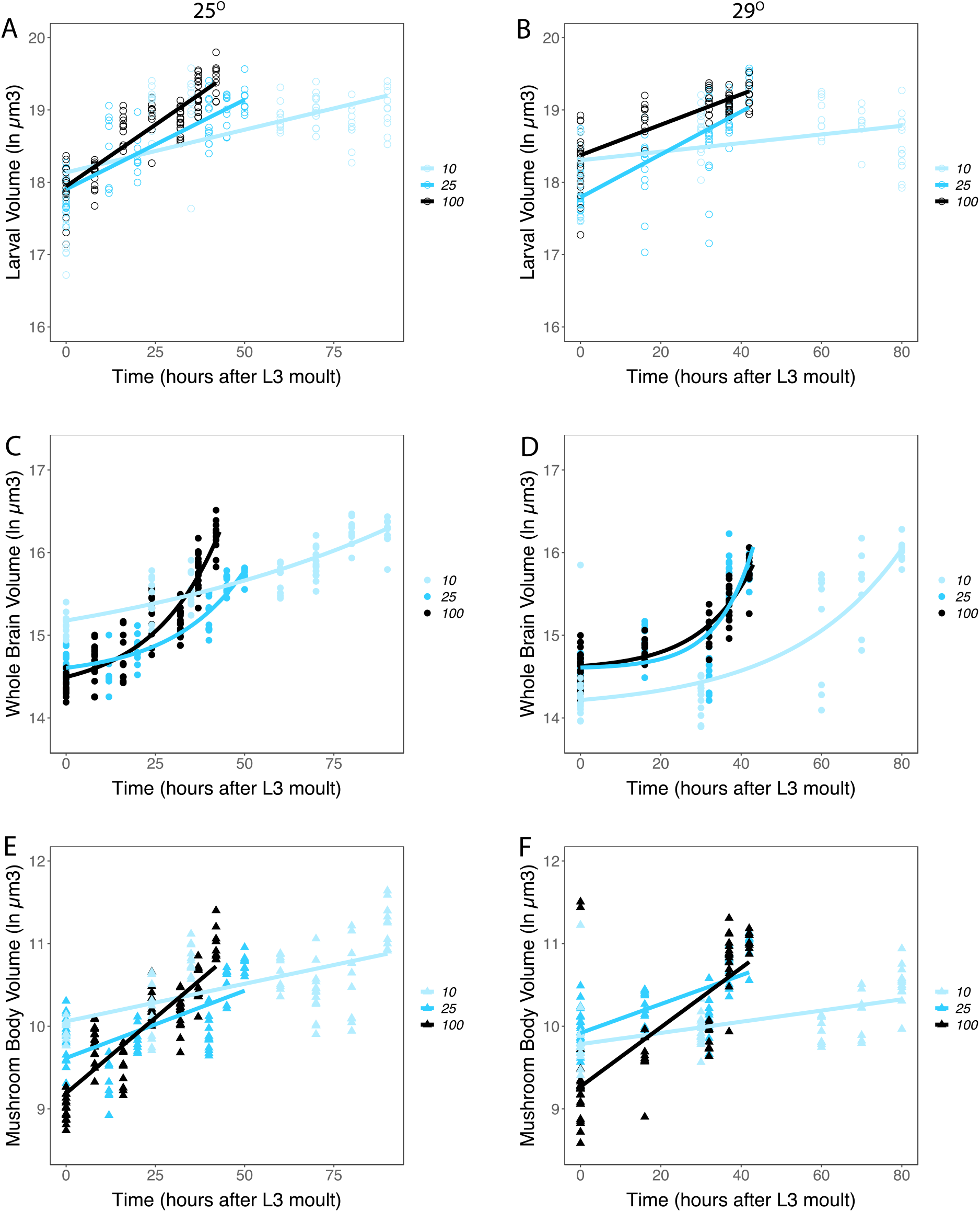
Growth rates of the larval body (A, B), whole brain (C, D), and mushroom bodies (E, F) over time from the moult prior to third instar through pupariation under different dietary and thermal conditions. Panels (A), C) and (E) show three dietary conditions (10%, 25%, and 100% food) at 25°C. Panels (B), (D) and (F) show the three dietary conditions (10%, 25%, and 100% food) at 29°C.

Changing developmental time allowed us to directly test our different models. We predicted that brain structures would remain robust to changes in developmental time in one of three ways (Figure 1A). Our first model predicted that when developmental time was extended, brain structures would maintain their growth rates, grow to their final size, and then stop growing and remain the same size until pupariation. This would be modelled best using an asymptotic regression, but could also be approximated by a negative quadratic term from a second order polynomial regression – indicating growth rates are slowing down. In our second model, we predicted that brain structures would remain robust against changes in developmental time by altering the time at which growth is initiated, but maintaining constant growth rates. This hypothesis would be best supported by a change in the lag constant of a lagged exponential regression. Our final hypothesis proposed that brain structures would carefully tune both their rates of growth and the time they initiated exponential growth, supported by a change in both the scaling and lag constants of a lagged exponential regression or by a change in slope in a linear regression in the case of the mushroom bodies.

In the mushroom body, we found that diluting the food reduced growth rates (Figure 2E, F, Supplementary Table 3), but that rearing temperature did not affect the rate of growth in this structure. This resulted in a significant decrease in growth rates for larvae grown on 10% food when compared to 25% food, as well as reduced growth rates on 25% food when compared to 100% food at both temperatures. Under all conditions, the mushroom bodies maintained linear growth trajectories. This best supports our model that at least the mushroom body compartment of the brain achieves robustness of size by carefully tuning its growth rates to adjust for changes in developmental time.

Because the whole brain showed non-linear growth patterns, we initially modelled whole brain growth using second order polynomials (Figure 2C, D, Supplementary Table 2). Similar to the larval body and mushroom bodies, diluting the food reduced the growth rates of the whole brain with the slowest growth on 10% food for both temperatures. Rearing temperature also reduced growth rates in the whole brain (Figure 2C, D, Supplementary Table 3), and the way that food affected growth rates depended on the rearing temperature. For larvae reared at 25ºC, growth rates differed depending on whether they were given 25% or 100% food. At 29ºC, there was no difference in growth rate between the 25% and 100% food. Thus, the whole brain shows complex responses to the combined effects of temperature and diet.

These models allowed us to further distinguish between our hypotheses. If whole brains grew to a target size and then stopped, we would expect the quadratic terms from our polynomial regressions to be negative as growth rates decreased. In all cases where the quadratic term was significant in our models, we found that the value was positive (Table 1). This suggests that our first model – that brains should grow to a target size then stop – is not supported by our data.

**Table 1:**
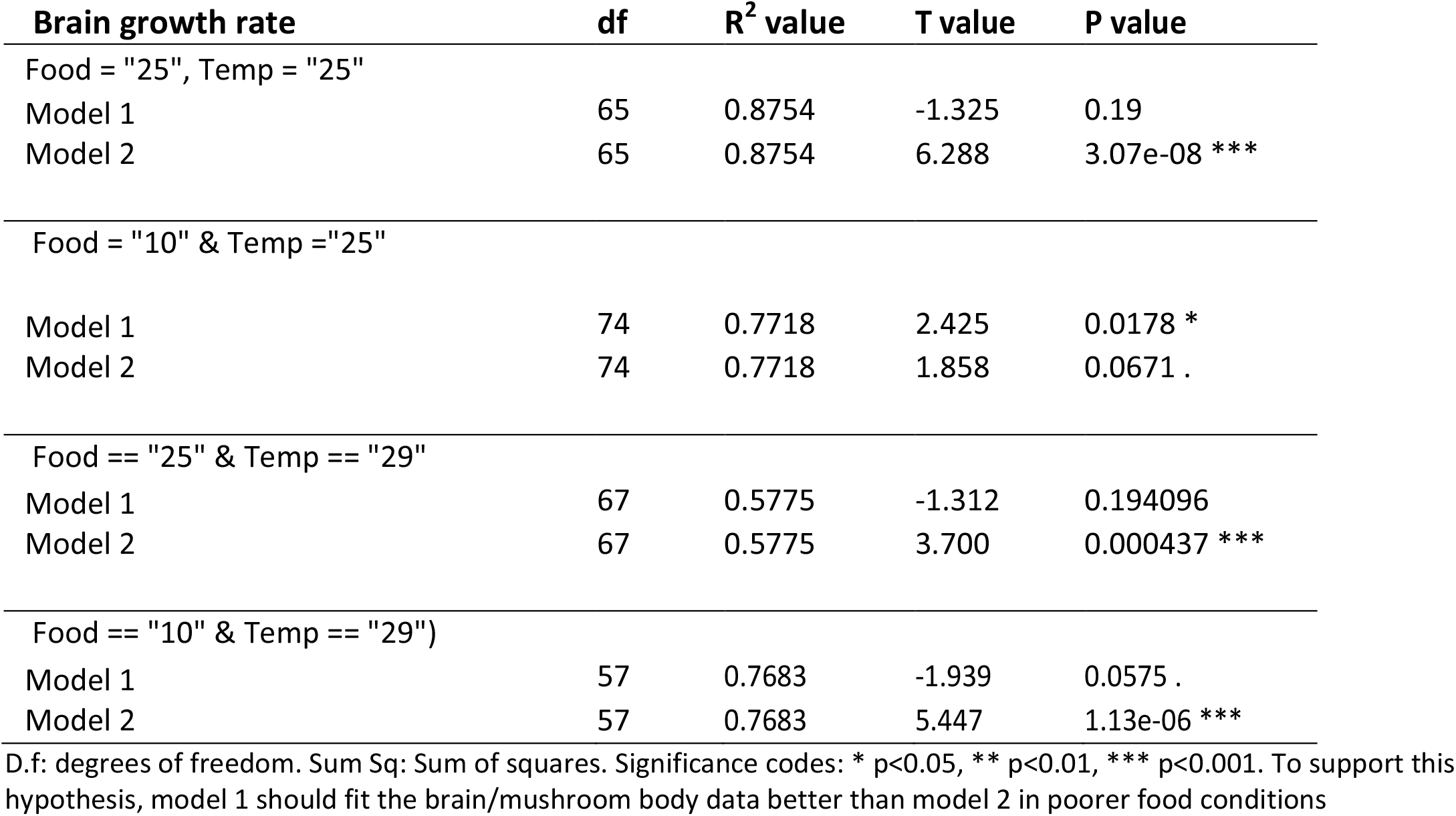
Model to test hypothesis 1 that brains maintain growth rate to target size when developmental time is extended

| Brain growth rate | df | R<sup>2</sup> value | T value | P value |
| --- | --- | --- | --- | --- |
| Food = "25", Temp = "25" |  |  |  |  |
| Model 1 | 65 | 0.8754 | -1.325 | 0.19 |
| Model 2 | 65 | 0.8754 | 6.288 | 3.07e-08 *** |
| Food = "10" & Temp ="25" |  |  |  |  |
| Model 1 | 74 | 0.7718 | 2.425 | 0.0178 * |
| Model 2 | 74 | 0.7718 | 1.858 | 0.0671 . |
| Food == "25" & Temp == "29" |  |  |  |  |
| Model 1 | 67 | 0.5775 | -1.312 | 0.194096 |
| Model 2 | 67 | 0.5775 | 3.700 | 0.000437 *** |
| Food == "10" & Temp == "29") |  |  |  |  |
| Model 1 | 57 | 0.7683 | -1.939 | 0.0575 . |
| Model 2 | 57 | 0.7683 | 5.447 | 1.13e-06 *** |

D.f: degrees of freedom. Sum Sq: Sum of squares. Significance codes: \* p<0.05, \*\* p<0.01, \*\*\* p<0.001. To support this hypothesis, model 1 should fit the brain/mushroom body data better than model 2 in poorer food conditions

We can distinguish between our second and third models using the lagged exponential growth models using the formula 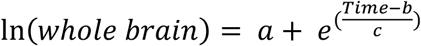, where a is the intercept, b is the lag constant, and c is the scaling constant. If brains remain robust to changes in developmental time by altering the time at which they turn on growth (hypothesis 2, Figure 1), we would expect the lag constant (b) to change, but not the scaling constant (c). Hypothesis 3 would be supported if both the lag constant (b) and scaling constant (c) changed with altered developmental time (Figure 1).

We fit our whole brain growth data with lagged exponential curves and explored whether the lag and scaling constants differed across our six environmental conditions (Table 2). We then conducted pairwise comparisons between whole brain growth curves either at the same temperature but across different diets, or on the same diet but across the two temperatures. We asked whether fitting specific lag and scaling constants for the curves for each condition improved the fit to the data. For the comparisons between the 10% food and either the 25% or the 100% food, the lag constants were too dissimilar to find a common coefficient, resulting in a failure to resolve a null model. While this suggests that the lag constants differ in these comparisons, we cannot formally test for this. However, both the lag constants (1 instance) and the scaling constants (5 instances) differed significantly between conditions for whole brain growth (Table 2). Taken together, our data best supports a model where both the timing at which exponential growth begins and the growth rate are carefully tuned to adjust for differences in developmental time.

**Table 2:** Model to test that brains remain robust to changes in developmental time by changing the time at which they turn on growth (hypothesis 2) or by changing both growth rates and the time at which they turned on growth changed (hypothesis 3).

| Comparison |  | Any constant differs | Lag or Scaling Constant Differ | Lag Constant Differs | Scaling Constant differs | Intercept Differs |
| --- | --- | --- | --- | --- | --- | --- |
| Temperature (°C) | Food (%) | F value (all constants the same) | (a varies) | (a and c varies) | (a and b varies) | (b & c varies) |
| 25 | 25 & 100 | 43.315*** | 48.136*** | 4.6947 | 0.0854 | - |
| 25 | 10 & 25 | does not resolve | does not resolve | does not resolve | 2.772*** | 3.4892*** |
| 25 | 10 & 100 | does not resolve | does not resolve | does not resolve | 3.673*** | 2.1648* |
| 29 | 25 & 100 | 0.8435 | - | - | - | - |
| 29 | 10 & 25 | 33.158*** | 16.125*** | does not resolve | 8.7448** | 4.9836 |
| 29 | 10 & 100 | 45.503*** | 14.132*** | does not resolve | 3.0147 | 2.8911 |
| 25 & 29 | 10 | 64.844*** | 29.453*** | 0.1876 | 5.2161* | - |
| 25 & 29 | 25 | 10.522*** | 13.738*** | 1.2351 | 7.3676** | - |
| 25 & 29 | 100 | 15.285*** | 15.828*** | 6.8819*** | 1.4653 | - |

D.f: degrees of freedom. Sum Sq: Sum of squares. Significance codes: \* p<0.05, \*\* p<0.01, \*\*\* p<0.001, ‘.’ p<0.1. To support hypothesis 2, the lag constant (b) should change, but not the scaling constant (c) and hypothesis 3, if both (b) and (c) changes with altered developmental time. We applied Holm’s adjustment to the p-values to account for multiple tests.

### Changing environmental conditions affects size traits in the prepupae

We have shown that the growth dynamics of the larval body, mushroom body, and whole brain are all sensitive to environmental perturbation, but that they respond in different ways to changes in diet and temperature. We next extended these findings by examining the effects of changed environmental conditions on their final size at pupariation.

Pupal body volume decreased as the food was diluted and also decreased at the higher temperature (Figure 3A, Table 3). This is what we would have expected given previously published data on the effects of diet and temperature on pupal body size (Couret et al., 2014; Davidowitz et al., 2003; Loeb & Northrop, 1917). At pupariation, we did not observe a significant effect of diet on its own for whole brain volume (Figure 3B, Table 3). However, whole brains were smaller at 29ºC than at 25ºC, and there was a significant temperature by diet interaction (Figure 3B, Table 3). This is due to the fact that at 25ºC larval diet had no effect on brain volume while at 29ºC, brain volume decreased with diet concentration. Mushroom body volumes at pupariation varied with diet and temperature, with increasing food concentrations and increasing temperatures negatively impacting mushroom body volume (Figure 3C, Table 3). The significant interaction between diet and temperature results from the fact that while food concentration correlates negatively with mushroom body volume at 25ºC, it correlates positively with mushroom body volume at 29ºC.

**Figure 3:**
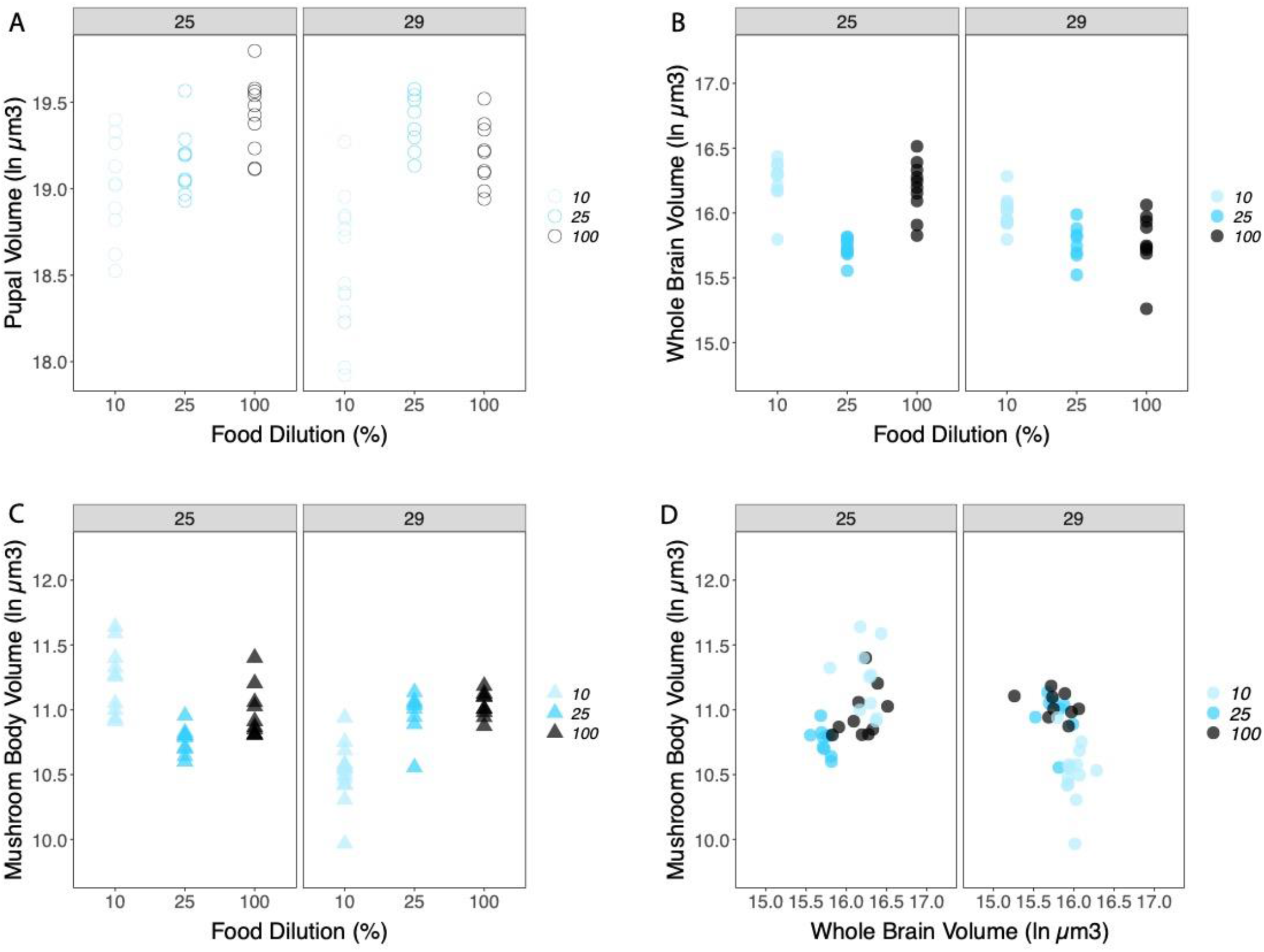
The prepupal volume of. (A), whole brain volume (B) and the mushroom body volume (C) across nutritional (10%, 25% and 100%) and thermal conditions (25°C and 29°C). The relationship between whole brain and mushroom body volume is shown in (D).

**Table 3:**
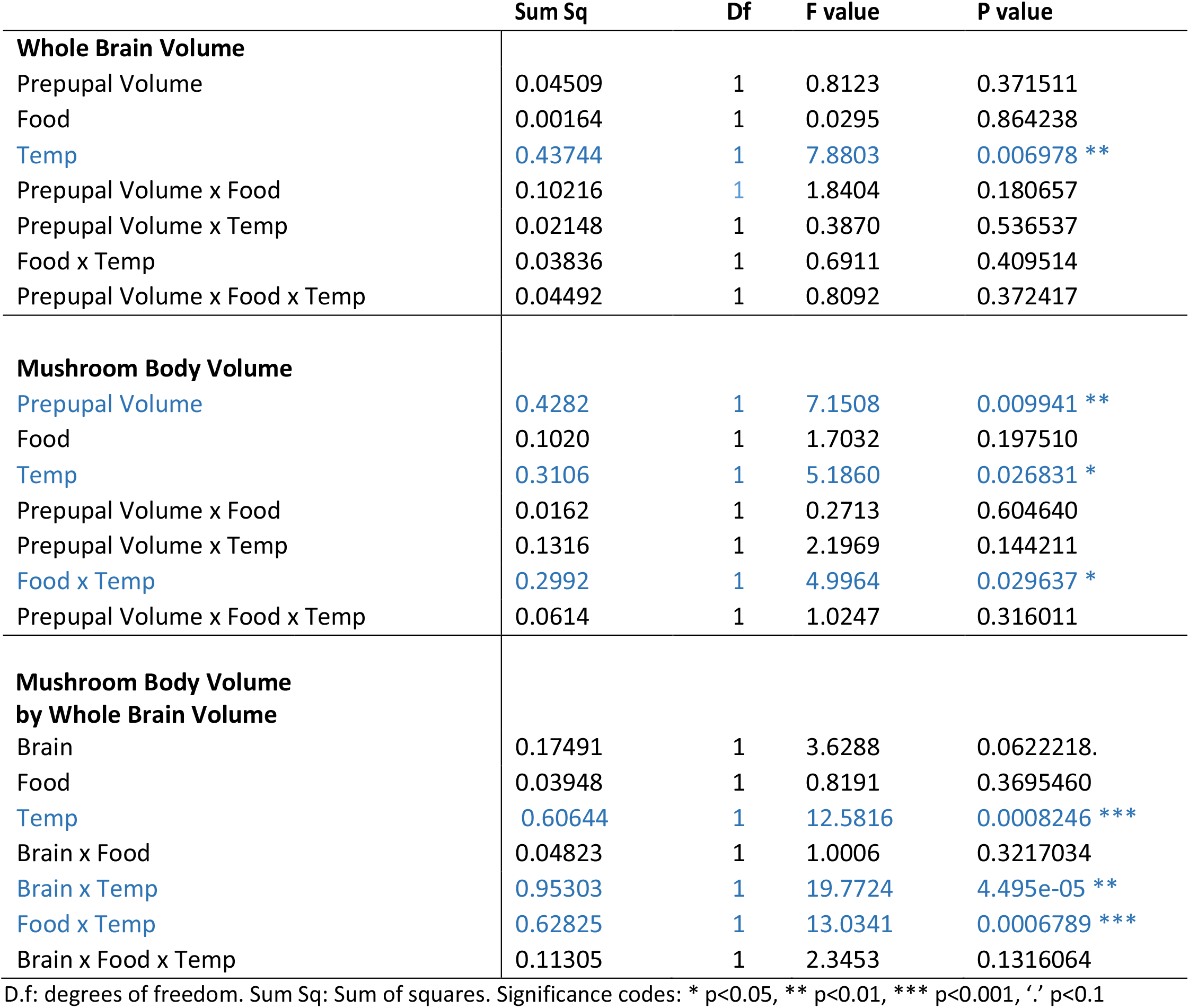
The final relationship between whole brain and body size depends only on temperature whereas the mushroom body/body size relationship depends on both diet and temperature, with a significant two – way interaction.

|  | Sum Sq | Df | F value | P value |
| --- | --- | --- | --- | --- |
| **Whole Brain Volume** |  |  |  |  |
| Prepupal Volume | 0.04509 | 1 | 0.8123 | 0.371511 |
| Food | 0.00164 | 1 | 0.0295 | 0.864238 |
| Temp | 0.43744 | 1 | 7.8803 | 0.006978 ** |
| Prepupal Volume x Food | 0.10216 | 1 | 1.8404 | 0.180657 |
| Prepupal Volume x Temp | 0.02148 | 1 | 0.3870 | 0.536537 |
| Food x Temp | 0.03836 | 1 | 0.6911 | 0.409514 |
| Prepupal Volume x Food x Temp | 0.04492 | 1 | 0.8092 | 0.372417 |
| **Mushroom Body Volume** |  |  |  |  |
| Prepupal Volume | 0.4282 | 1 | 7.1508 | 0.009941 ** |
| Food | 0.1020 | 1 | 1.7032 | 0.197510 |
| Temp | 0.3106 | 1 | 5.1860 | 0.026831 * |
| Prepupal Volume x Food | 0.0162 | 1 | 0.2713 | 0.604640 |
| Prepupal Volume x Temp | 0.1316 | 1 | 2.1969 | 0.144211 |
| Food x Temp | 0.2992 | 1 | 4.9964 | 0.029637 * |
| Prepupal Volume x Food x Temp | 0.0614 | 1 | 1.0247 | 0.316011 |
| **Mushroom Body Volume by Whole Brain Volume** |  |  |  |  |
| Brain | 0.17491 | 1 | 3.6288 | 0.0622218. |
| Food | 0.03948 | 1 | 0.8191 | 0.3695460 |
| Temp | 0.60644 | 1 | 12.5816 | 0.0008246 *** |
| Brain x Food | 0.04823 | 1 | 1.0006 | 0.3217034 |
| Brain x Temp | 0.95303 | 1 | 19.7724 | 4.495e-05 ** |
| Food x Temp | 0.62825 | 1 | 13.0341 | 0.0006789 *** |
| Brain x Food x Temp | 0.11305 | 1 | 2.3453 | 0.1316064 |

D.f: degrees of freedom. Sum Sq: Sum of squares. Significance codes: \* p<0.05, \*\* p<0.01, \*\*\* p<0.001, '.' p<0.1

These differences in the way the whole brain and mushroom body volumes respond to diet and temperature has interesting implications for brain shape. While mushroom body volumes are remarkably robust in size on 25% and 100% foods, on 10% food they are larger for their brain size at 25ºC and smaller for their whole brain size at 29ºC (Figure 3D, Table 3). This highlights that brain shape changed across environmental conditions, as compartments of the brain differed in how they grew in response to these conditions.

## Discussion

Individual organs vary in their response to the environmental conditions that affect adult body size (Shingleton et al., 2009). Organs like the brain and genital discs are known to be nutritionally insensitive when compared to organs like the wing (Chell & Brand, 2010; Cheng et al., 2011; Shingleton, 2010; Shingleton & Frankino, 2018; Tang et al., 2011). While we have some understanding of the genetic mechanisms underpinning robustness in size in these organs, our understanding of how these mechanisms affect the dynamics of growth was poorly understood. Further, the brain is a complex organ whose compartments do not all behave in the same manner. Functional compartments like the *Drosophila* mushroom body differ in their growth patterns as well as their nutritional plasticity from the rest of the brain. In this study, we aimed to test our predictions that the differences in proliferation between neurons of the whole brain and mushroom bodies would confer distinct growth dynamics, which could impart differences in their sensitivity to environmental conditions.

Previous studies had suggested that brain sparing occurs under stressful conditions because Jeb/Alk maintain high levels of insulin and TOR signalling in the neuroblasts (Cheng et al., 2011). These same conditions act to extend development time of the larva (Beadle et al., 1938; Nunney & Cheung, 1997; Partridge et al., 1994; Robertson, 1962, 1966; Robertson & Reeve, 1952). If insulin and TOR levels are at the same level in the brains of starved and fed larvae, then the brain must have additional mechanisms to prevent overgrowth when development time is extended. In this study, we altered development time by changing both nutrition and temperature. We proposed three hypotheses that would allow brain size to remain robust against environmental conditions. These posited that in response to changes in total development time the brain would either 1) grow to a target size and stop growing for the remainder of the growth period, 2) delay the onset of its growth, but maintain constant growth rates even under stressful conditions, or 3) regulate both its growth rate and the time at which it switches growth on to adjust for changes in developmental time. Our data supports our third hypothesis, that robustness of brain size is possible because both the time at which exponential growth is initiated and the rates of growth of the brain have been altered.

Our results imply that Jeb/Alk signalling, which is responsible for brain sparing in *Drosophila*, plays a more nuanced role than previously described. Rather than simply maintaining high levels of insulin and/or TOR signalling, signalling from Alk could be acting to adjust growth rates of the brain to match changes in developmental time. Precisely how this occurs is unknown, however given that both insulin and ecdysone signalling are key regulators of the length of the growth period (Caldwell et al., 2005; Colombani et al., 2005; Koyama et al., 2014; Mirth et al., 2005; Shingleton et al., 2005), these systemic cues could be regulating the concentration of Jeb secreted by the glial cells in accordance with the degree to which development is delayed. Other organs that show robustness in final size could be responding to environmental conditions in a similar fashion. For example, we would predict the genital discs maintain robust final size by tuning their growth rates to account for extended growth periods under poor nutrition or thermal stress.

While the size of the pupal brain is robust against environmental conditions, this does not mean that brain growth is insensitive to environmental stress. Nutritional signals are important for neuroblasts to exit quiescence and re-initiate proliferation in the larval stages (Britton & Edgar, 1998; Chell & Brand, 2010; Yuan et al., 2020). Cues from the fat body, the insect equivalent of the adipose tissue and liver, signal to glial cells in the brain, which in turn produce insulin-like peptides that induce the neuroblasts to recommence cell divisions (Britton & Edgar, 1998; Chell & Brand, 2010; Yuan et al., 2020). Starving larvae in early instars causes most neuroblasts and glia, with the exception of the mushroom body neuroblasts, to remain quiescent (Britton & Edgar, 1998; Chell & Brand, 2010; Yuan et al., 2020). This is owing to the cell-autonomous and non-autonomous growth coordination activity of PI3Kinase in the early larval stages of development (Yuan et al., 2020). After they exit quiescence, neuroblasts no longer depend on nutritional cues to maintain proliferation (Cheng et al., 2011; Lanet & Maurange, 2014). However, our data demonstrates that rates of brain growth remain sensitive to environmental cues. Whether this is due to changes in rates of neuroblast proliferation, or changes in the rates of increase in cell size within the brain is yet unclear.

Although the growth dynamics of the whole brain change to accommodate additional developmental time, our findings also demonstrate that not all compartments of the brain should be expected to respond in the same way. Our comparisons between whole brain and the mushroom bodies highlight how the growth dynamics of specific brain compartments can differ from the patterns observed across the brain as a whole. Some of these differences arise simply due to differences in the timing of neuroblast reactivation. While the neurons of the mushroom body continue to proliferate throughout larval development, most other neuroblasts reinitiate proliferation after the late second instar. This alone should be sufficient to generate differences in growth dynamics between the mushroom bodies and the rest of the brain, however the role of increases in cell size across brain regions has yet to be explored.

Furthermore, differences in growth patterns are not unique to the mushroom body. Unlike most of the other regions of the brain, the optic lobe shows extensive plasticity in size with nutritional conditions (Lanet et al., 2013; Lanet & Maurange, 2014). This is presumably to compensate for changes in eye size across environmental conditions, and is facilitated by their unique mode of development. Instead of arising from embryonic neuroblasts, the optic lobe is formed from neuroepithelium that continues to divide and expand until early in the third instar (Brand & Livesey, 2011; Farkas & Huttner, 2008). Proliferation of the optic lobe neuroepithelium remains sensitive to nutrition until the early third instar, where a small pulse of ecdysone induces the cells in this neuroepithelium to become neuroblasts (Lanet et al., 2013). After this transition, starvation no longer impacts cell divisions in this brain region, and each neuroblast proceeds to divide and generate the full complement of neuronal cells types necessary for the function of the optic lobe (Lanet et al., 2013). This ensures that while the total number of neurons in the optic lobe is plastic, the diversity of cell types is held constant (Lanet et al., 2013). Given its mode of development and persistent sensitivity to nutrition, we would expect that the optic lobes would also exhibit different growth dynamics from the whole brain.

Given these differences in growth patterns across the mushroom body and whole brain, we would predict that the compartments of the brain might differ in their sensitivity to the two pathways known to regulate growth in response to nutrition: the insulin and TOR pathways (Yuan et al., 2020). Other studies of whole brain growth in *Drosophila* (Sousa-Nunes et al., 2010; Yuan et al., 2020), and in mammals (Cloetta et al., 2013), show TOR signalling controls cell cycle progression and neuronal exit from quiescence respectively, ultimately regulating final brain size. In the mushroom body, the Pax-6 orthologue, Eyeless, allows mushroom body neuroblasts to continue proliferating independent of PI3Kinase activity, a central regulator of insulin signalling, under conditions of poor nutrition (Sipe & Siegrist, 2017). This is likely to be a matter of degree: while eyeless undoubted plays a role in maintaining proliferation, insulin signalling in the mushroom body neuroblasts has its own independent effects on proliferation and in controlling the size of the arbour (Sipe & Siegrist, 2017).

Finally, the majority of studies of brain growth have focused on nutritional stress. However, a number of other conditions are known to extend developmental time, including temperature, oxygen limitation, and larval density (Mirth & Shingleton, 2012; Partridge et al., 1994; Peck & Maddrell, 2005). The mechanisms regulating extended developmental time under these conditions are less well understood, but ultimately culminate in changing the rate of ecdysone production and secretion. Previous studies have documented that reducing or eliminating ecdysone or ecdysone signalling also reduces brain size (Herboso et al., 2015; Lanet et al., 2013). Thus, in addition to insulin and TOR pathways, ecdysone is likely to regulate the size of whole brains and the size of its compartments by fixing the length of their growth period.

## Conclusion

In this research, we sought to understand how organs achieve robust final size by exploring the growth dynamics of the brain across nutritional and thermal conditions. We found that at least one compartment of the brain can differ in its growth patterns from the rest of the brain, and speculate that this might be true of other compartments. These distinct growth patterns allow specific brain regions to vary their response to changing environmental conditions. Taken together, our findings demonstrate that brain compartments achieve robustness in final size via different trajectories. Furthermore, by probing the growth dynamics of organs under environmental stress, we fill in important gaps in our knowledge of how these organs achieve robustness of final size.

## Acknowledgements

We are grateful to Keith Schulze, Alex Fulcher and Stephen Firth of the Monash MicroImaging (MMI) for their technical support and members of the Mirth lab for their helpful discussion on the project. This work was supported by an Australian Research Council Future Fellowship (FT170100259) to CKM and the School of Biological Sciences, Monash University, Australia.

## Author contributions

AEC, BN and CKM conceived and designed the experiments. AEC performed the experiments. AEC and CKM analysed the data. All authors contributed to interpretation of data and final manuscript preparation.

## Declaration of interests

The authors declare no competing interests.

## SUPPLEMENTARY MATERTIAL

**Supplementary Figure 1:**
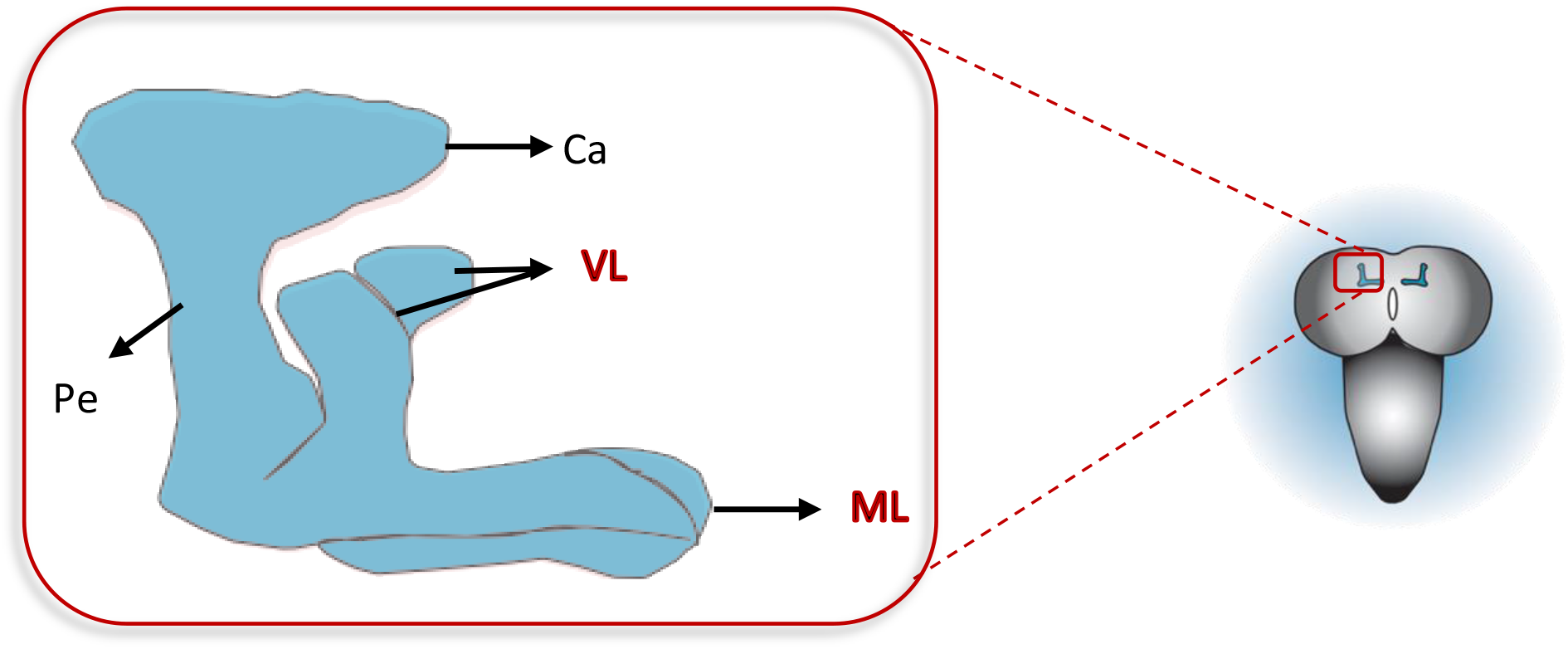
Schematic of the dorsal view of mushroom body neuropil in the left hemisphere of brain of *Drosophila melanogaster* showing the Calyx (Ca), Peduncle (Pe), Vertical lobe (VL) and Medial lobe. Regions included in this study are labelled in red, the vertical and medial lobes only.

**Supplementary Figure 2a:**
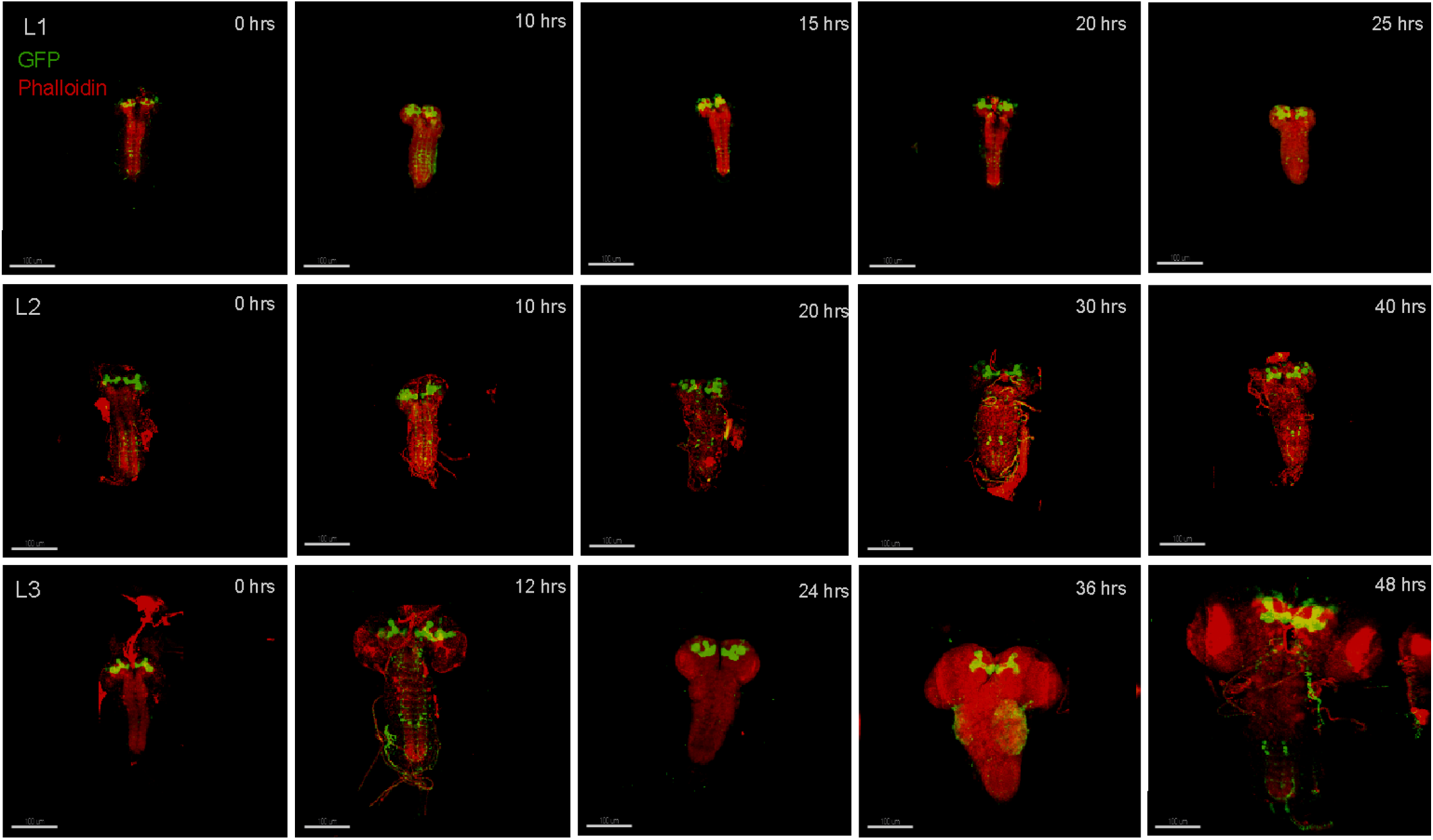
Changes in brain growth across larval stage of development. Larval brains expressing GFP in the neurons (green) of the mushroom body co-stained with phalloidin (red) across five developmental time points in the three larval stages. First instar (L1) A-E (0 hrs is relative to hatching), the second instar (L2) F-J (0 hours relative to the moult to L2) and the third instar (L3) K-P (0 hours relative to the moult to L3). At L3, the last two time points correspond to wandering and white prepupal stages. (Scale bar: 100μm)

**Supplementary Figure 2b:**
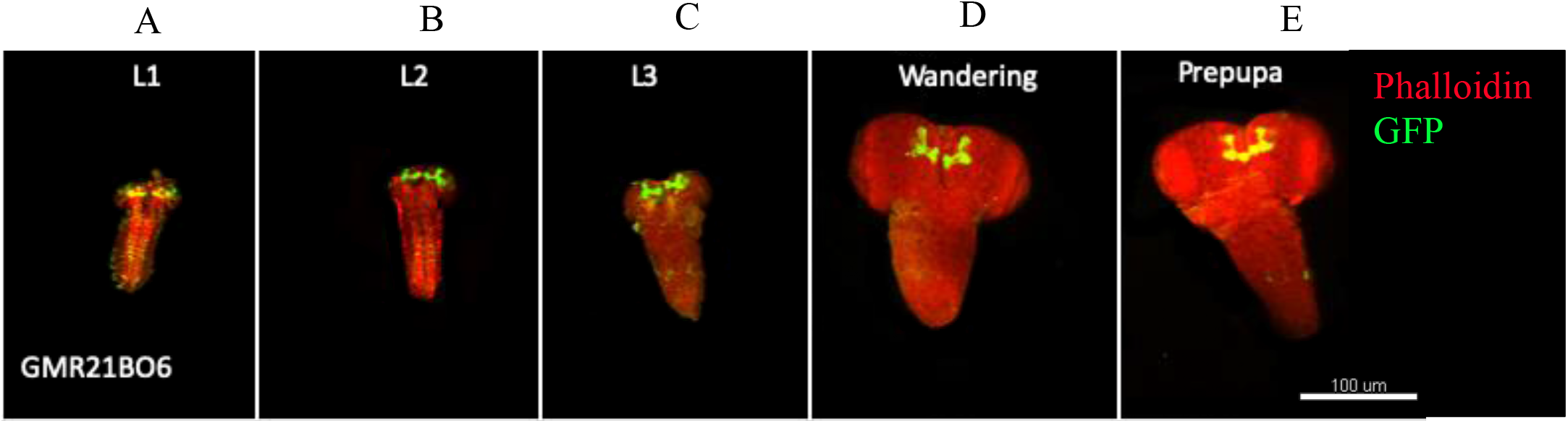
Larval brains expressing GFP in the neurons (green) of the mushroom body co-stained with phalloidin (red) across five different stages. (a-e) shows brains at 0hr of L1, L2, L3, wandering and white prepupae larval stages respectively.

**Supplementary Figure 3:**
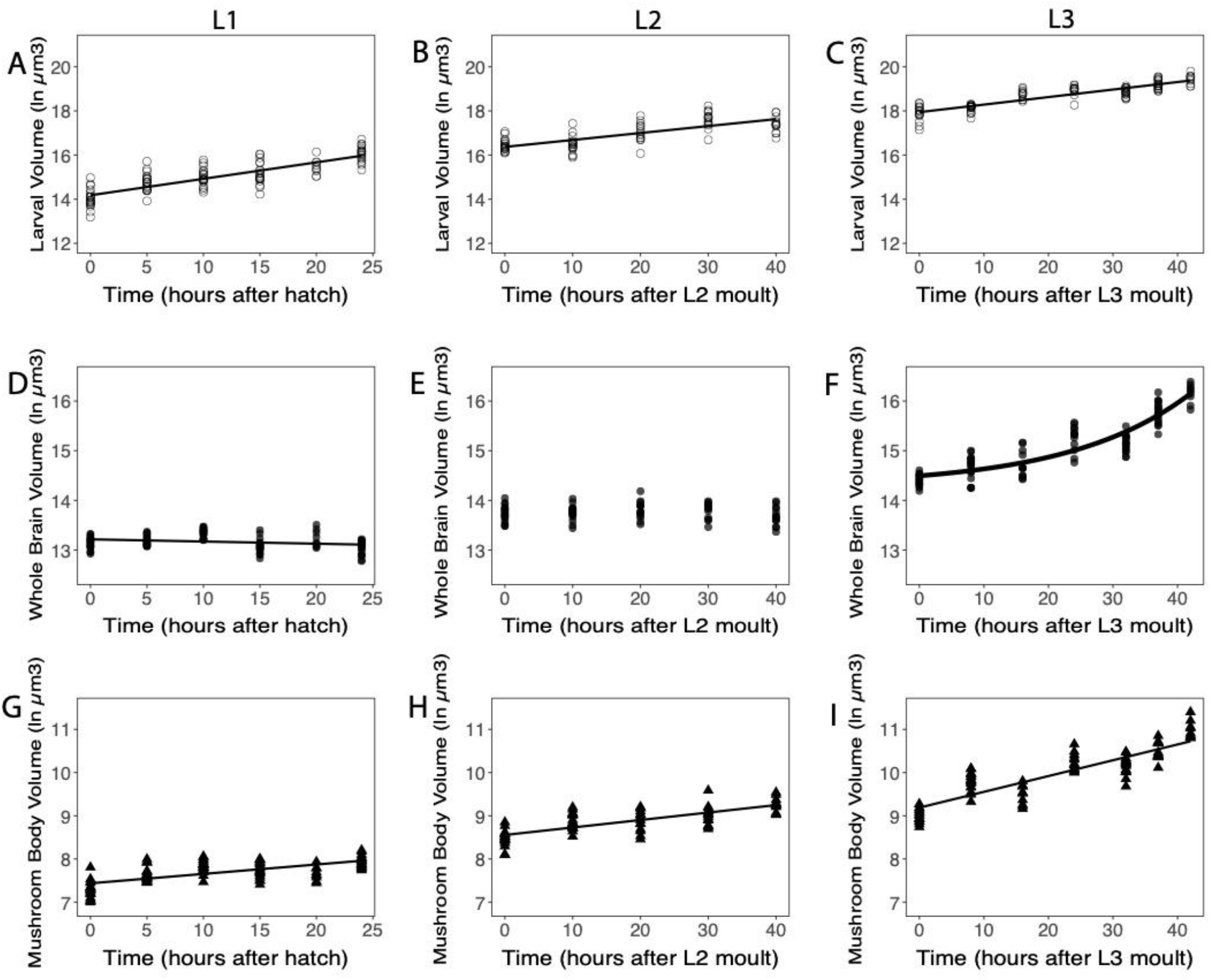
Growth patterns of larval body, brain and mushroom body. The volume of the larval body (A-C), whole brain (D-F), and mushroom body (G-I) at L1 stage (A, D, G), L2 stage (B, E, H) and at L3 stage (C, F, I) measured from 0 hours after hatching/ larval moult to the end of the larval instar. At L3, the last two timepoints correspond to wandering and white prepupae larval stages. Each point shows individuals measured.

**Supplementary Figure 4:**
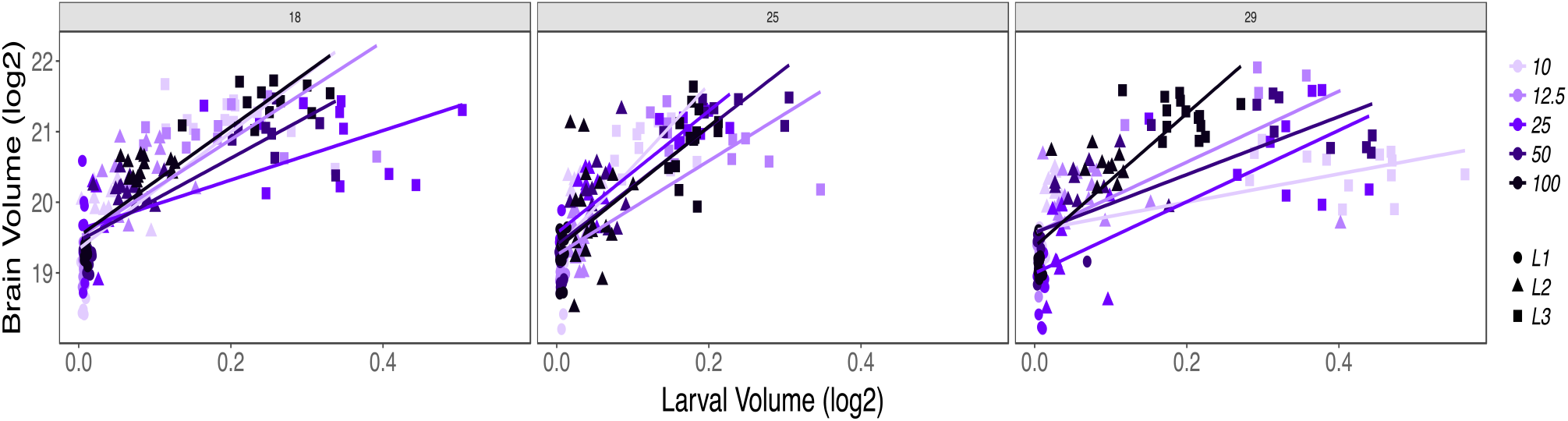
showing the effect of temperature and nutrition on the brain volume across development. Each box represents the different temperatures, and the concentration of food is represented in the different colour codes where the highest food concentration is shown in black and the lowest food concentration is seen in lilac colour.

**Supplementary Table 1:** Linear regression models of larval volume, brain and mushroom body volume across the first, second and third instar stages of development.

| Trait | Stage | F stat | p value | R<sup>2</sup> Adj |
| --- | --- | --- | --- | --- |
| Larval Volume | L1 | 224.44 | < 2.2e-16 | 0.6762 |
|  | L2 | 99.333 | 1.28E-15 | 0.5514 |
|  | L3 | 374.65 | < 2.2e-16 | 0.7806 |
| Brain Volume | L1 | 6.1501 | 0.01472 | 0.04592 |
|  | L2 | 0.0045 | 0.9468 | -0.0126 |
|  | L3 | 372.81 | < 2.2e-16 | 0.7798 |
| MB Volume | L1 | 83.835 | 4.45E-15 | 0.4364 |
|  | L2 | 99.097 | 1.35E-15 | 0.5508 |
|  | L3 | 354.69 | < 2.2e-16 | 0.7711 |

R<sup>2</sup> Adj: Adjusted R<sup>2</sup>. Significance codes: \* p<0.05, \*\* p<0.01, \*\*\* p<0.001, '.' p<0.1

**Supplementary Table 2:** Akaike’s Information Criterion (AIC) and Bayesian Information Criteria (BIC) for modelling larval volume, brain and mushroom body volume in the third instar (L3) stage of development. Values for best fit are in blue.

| Trait | Fit | AIC | BIC |
| --- | --- | --- | --- |
| Body | Volume.lmL3 | 20.35881 | 28.34913 |
|  | Volume.lmL3poly | 19.63951 | 30.29327 |
|  | Volume.expL3 | 30.84764 |  |
|  | Volume.explagL3_100 |  | 38.83796 |
|  | Volume.powerL3 | 118.6141 | 126.6044 |
| Brain | Brain.lmL3 | 36.96086 | 44.95118 |
|  | Brain.lmL3poly | 18.70613 | 29.35988 |
|  | Brain.expL3 | 19.51189 | 27.5022 |
|  | Brain.explagL3_100 | 12.13046 | 22.78422 |
|  | Brain.powerL3 | 156.5104 | 164.5007 |
| Mushroom Body | MB.lmL3 | 40.93267 | 48.92299 |
|  | MB.lmL3poly | 42.85683 | 53.51059 |
|  | MB.expL3 | 42.70775 | 50.69807 |
|  | MB.explagL3_100 | 42.80673 | 53.46049 |

lm: linear model, poly: polynomial, exp: exponential model, explag: lagged exponential model.

**Supplementary Table 3:** Growth rates of the larval body, brain and mushroom bodies depend on nutritional and thermal conditions. Larval body and mushroom body volumes were fit with linear models (lm). Brain volumes were fit with second order polynomial models with Time as (Time, 2, raw = TRUE).

| Larval Volume | Sum Sq | Df | F value | P value |
| --- | --- | --- | --- | --- |
| Time | 58.358 | 1 | 402.9612 | <2.2e-16 *** |
| Food | 19.295 | 2 | 66.6153 | < 2.2e-16 *** |
| Temp | 0.051 | 1 | 0.3500 | 0.554400 |
| Time x Food | 16.805 | 2 | 58.0189 | < 2.2e-16 *** |
| Time x Temp | 1.358 | 1 | 9.3796 | 0.002328 ** |
| Food x Temp | 0.429 | 2 | 1.4828 | 0.228125 |
| Time x Food x Temp | 1.755 | 2 | 6.0605 | 0.002532 ** |
| **Brain Volume** |  |  |  |  |
| (Time, 2, raw = TRUE) | 89.118 | 2 | 601.0508 | < 2.2e-16 *** |
| Food | 2.620 | 2 | 17.6721 | 4.178e-08 *** |
| Temp | 5.342 | 1 | 72.0532 | 3.331e-16 *** |
| (Time, 2, raw = TRUE) x Food | 13.364 | 4 | 45.0669 | < 2.2e-16 *** |
| (Time, 2, raw = TRUE) x Temp | 6.363 | 2 | 42.9132 | < 2.2e-16 *** |
| Food x Temp | 14.659 | 2 | 98.8697 | < 2.2e-16 *** |
| (Time, 2, raw = TRUE) x Food x Temp | 2.207 | 4 | 7.4441 | 8.318e-06 *** |
| **Mushroom Body Volume** |  |  |  |  |
| Time | 51.040 | 1 | 371.8073 | < 2.2e-16 *** |
| Food | 2.806 | 2 | 10.2220 | 4.576e-05 *** |
| Temp | 0.012 | 1 | 0.0880 | 0.7669 |
| Time x Food | 25.315 | 2 | 92.2044 | < 2.2e-16 *** |
| Time x Temp | 0.081 | 1 | 0.5931 | 0.4416 |
| Food x Temp | 6.580 | 2 | 23.9649 | 1.318e-10 *** |
| Time x Food x Temp | 0.104 | 2 | 0.3788 | 0.6849 |

D.f: degrees of freedom. Sum Sq: Sum of squares. Significance codes: \* p<0.05, \*\* p<0.01, \*\*\* p<0.001, ' ' p<0.1

## Materials and Methods

### Fly strains and husbandry

*Drosophila* stocks were reared at 25°C with 65% humidity, on a 12-hour light/dark cycle and maintained on sucrose-yeast (SY) diet (detailed below). To drive the expression of green fluorescent protein (GFP) in the mushroom body neurons, we used the R21B06-GAL4 line (BDSC 68318), known to be expressed in the mushroom bodies of larval and adult brains (http://flweb.janelia.org/cgi-bin/flew.cgi; (Jenett et al., 2012; Pfeiffer et al., 2008). This line was crossed with a membrane-tagged GFP reporter (w[*]; P[y[+t7.7] w[+mC]=10XUAS-IVS-myr::GFP]attP2). These stocks were obtained from the Bloomington Drosophila Stock Center, Indiana University, Bloomington.

### Media and larval rearing and staging conditions

Sucrose/Yeast (SY) diet was prepared as reported by (Toivonen et al., 2007), with 100g autolyzed Brewer’s Yeast, 50 g sucrose, 10 g agar, 1.5 ml propionic acid, 15 ml Nipagin M solution dissolved in 1 L of distilled water. In addition to the standard diet (100% SY), we exposed larvae to additional experimental diets, which contained 10% and 25% of the caloric content of the standard SY diet. These diets were made by adding appropriate amounts of the original Brewer’s yeast and sucrose to the same concentration of agar and water. 25% food contained 25 g autolyzed Brewer’s Yeast and 12.5 g sucrose, while 10% food contained 10 g autolyzed Brewer’s Yeast and 5 g sucrose, dissolved in 1 L of distilled water. All diets were allowed to cool to 60° before the preservatives (propionic acid and Nipagin M) were added and the food dispensed.

Egg collection was carried out on normal diet without additional yeast for age synchronization. 100-150 eggs were transferred to a 60 × 15mm petri dish to control for population density. Newly-hatched first instar larva were collected in two hour cohorts starting 24 h after egg lay. These newly-hatched larvae were then staged to the appropriate time before imaging for body size measurements and dissection. To collect staged L2 and L3 larvae, we collected newly-moulted second and third larval stages, determined by their anterior spiracle morphology, in 2 hour cohorts as in (Mirth et al., 2005). These larvae were then staged to the desired time before imaging and dissection. To determine the duration from third instar to the white prepupal stage, L3 larvae were observed every 8 hours until all larvae pupariated. We defined pupariation as cessation of movement with evaginated spiracles (Koyama et al., 2014). Animals were raised at a control temperature of 25°C and experimental temperature of 29°C. All experiments were performed in three replicates on a 12 hr:12 hr light:dark cycle at 65% humidity.

### Image analysis and brain size measurement

Z-stack images were obtained from brain samples using the Leica Sp8 confocal microscope and 3-Dimensional volume was reconstructed with the Imaris© (Bitplane) software. Image normalization was performed to ensure standardized measurements across images with different signal intensities, and 3D analysis of the brain was done by software’s in-built wizard. Images were rendered, and brain size measurement was gotten as 3D volumes using the surface analysis tool on Imaris.

### Body size measurement, organ dissection, and immunocytochemistry

Animals picked at the relevant time points were first placed in cold PBS solution, to immobilize them, and then imaged using a Zeiss Stemi 508 dissecting microscope before dissection. These images were analyzed using the FIJI (ImageJ, Version 2.0.0-rc-69/1.52i, 2019) software. The length and width of the larva or pupa was measured using the straight-line tool, and volume was calculated using the formula *lw*^*2*^ (length x width^2^).

After measuring each larva/pupa, we dissected out their brain in cold 1x Phosphate Buffered Saline (1xPBS) under a Leica S9E dissecting microscope according to methods previously described (Daul et al., 2010; Hafer & Schedl, 2006). Isolated brains were fixed overnight in 4% paraformaldehyde at 4°C. After four washes in a solution of cold 0.3% Triton X-100 in PBS (PBT) for 20 minutes each, samples were incubated in 2% normal donkey serum block solution prior to immunostaining. The blocked tissue samples were then transferred to Acti-stain ™ 670 Phalloidin (1:1200, Cytoskeleton Cat#: PHDN1) reagent diluted in PBT and normal donkey serum, and incubated on a rocking platform shaker in the dark for 2-3days at 4°C. Prior to imaging, samples were rinsed in cold PBS, and PBS was replaced with 25% KY jelly in water solution. Samples were imaged using the Leica SP8 HyD microscope with 40x water immersion objective.

### Image processing and statistical analysis

Data analyses were carried out in R Studio (Version 1.2.5019^©^ 2009-2019 RStudio, Inc.). We fit our body and organ size data with both linear, using the *lm* function, and non-linear models, using the *nls* package (Baty et al., 2015). We used AIC and BIC to assess model fit, selecting the simplest models when these values were similar. Data visualization was conducted using the ggplot package (Wickham, 2016) in R Studio (Version 1.2.5019^©^ 2009-2019 RStudio, Inc.).

## Notes

### Competing Interest Statement

The authors have declared no competing interest.

